# Rapid evolution of distinct *Helicobacter pylori* subpopulations in the Americas

**DOI:** 10.1101/069070

**Authors:** Kaisa Thorell, Koji Yahara, Elvire Berthenet, Daniel J. Lawson, Ikuko Kato, Alfonso Mendez, Federico Canzian, María Mercedes Bravo, Rumiko Suzuki, Yoshio Yamaoka, Javier Torres, Samuel K Sheppard, Daniel Falush

**Affiliations:** Dept. of Microbiology, Tumour and Cell Biology, Karolinska Institutet, Sweden; Dept. of Bacteriology II, National Institute of Infectious Diseases, Japan; Medical Microbiology and Infectious Disease group, Swansea University, Wales; Integrative Epidemiology Unit, School of Social and Community Medicine, University of Bristol, UK; Karmanos Cancer Institute, Wayne State University, United States; Escuela Nacional de Ciencias Biologicas, IPN, Mexico; German Cancer Center, Heidelberg; Grupo de Investigación en Biología del Cáncer, Instituto Nacional de Cancerología, Colombia; Dept. of Environmental and Preventive Medicine, Oita University Faculty of Medicine, Japan; Unidad de Investigación en Enfermedades Infecciosas, UMAE Pediatria, IMSS, Mexico; Milner Center for Evolution, Dept. of Biology and Biochemistry, University of Bath, UK

**Keywords:** population structure, admixture, selection, chromosome painting, *Helicobacter pylori*

## Abstract

For the last 500 years, the Americas have been a melting pot both for genetically diverse humans and for the pathogenic and commensal organisms associated with them. One such organism is the stomach dwelling bacterium *Helicobacter pylori*, which is highly prevalent in Latin America where it is a major current public health challenge because of its strong association with gastric cancer. By analyzing the genome sequence of *H. pylori* isolated in North, Central and South America, we found evidence for admixture between *H. pylori* of European and African origin throughout the Americas, without substantial input from pre-Columbian (hspAmerind) bacteria. In the US, strains of African and European origin have remained genetically distinct, while in Colombia and Nicaragua, bottlenecks and rampant genetic exchange amongst isolates have led to the formation of national gene pools. We found four outer membrane proteins with atypical levels of Asian ancestry in American strains, including the adhesion factor AlpB, suggesting a role for the ethnic makeup of hosts in the colonization of incoming strains. Our results show that new *H. pylori* subpopulations can rapidly arise, spread and adapt during times of demographic flux, and suggest that differences in transmission ecology between high and low prevalence areas may substantially affect the composition of bacterial populations.

**Author Summary:** *Helicobacter pylori* is one of the best studied examples of an intimate association between bacteria and humans, due to its ability to colonize the stomach for decades and to transmit from generation to generation. A number of studies have sought to link diversity in *H. pylori* to human migrations but there are some discordant signals such as an “out of Africa” dispersal within the last few thousand years that has left a much stronger signal in bacterial genomes than in human ones. In order to understand how such discrepancies arise, we have investigated the evolution of *H. pylori* during the recent colonization of the Americas. We find that bacterial populations evolve quickly and can spread rapidly to people of different ethnicities. Distinct new bacterial subpopulations have formed in Colombia from a European source and in Nicaragua and the US from African sources. Genetic exchange between bacterial populations is rampant within Central and South America but is uncommon within North America, which may reflect differences in prevalence. Our results also suggest that adaptation of bacteria to particular human ethnic groups may be confined to a handful of genes involved in interaction with the immune system.

## Introduction

In 1492, Christopher Columbus initiated a rapid colonization of the New World, principally by European migrants and Africans brought as slaves that had catastrophic consequences for the indigenous population. The new migrants brought unfamiliar weapons and pathogens, including new populations of the stomach-colonizing bacterium *Helicobacter pylori. H. pylori* can persist for decades in the stomach, and is often transmitted vertically from parent to child but can also be acquired from individuals in close proximity. *H. pylori* evolves rapidly by both mutation and homologous recombination with other co-colonizing strains [1].

Studies of the global diversity of *H. pylori* have shown that Europeans, Africans and Native Americans carry genetically distinct populations of bacteria; named hpEurope, hpAfrica1 and hpAfrica2, and hspAmerind, respectively [2]. The relationships between bacterial populations reflect differentiation that has arisen during the complex migration history of humans, with the prefix “hp” indicating a population and “hsp” indicates a subpopulation, which are genetically distinct from each other but less differentiated than populations. hspAmerind bacteria are presumed to be descendants of the strains present in the Americas prior to 1492, and are a subpopulation of hpEAsia, which is found in Asian countries such as China and Japan. However, these strains are rare even within groups with substantial Native American ancestry and may being dying out in competition with other strains, due to low diversity within the population or other factors [3]. hpEurope bacteria are themselves ancient hybrids between two populations, whose close relatives are currently found in unadmixed populations in North East Africa (hpNEAfrica) and central Asia (hpAsia2). The Tyrolean Iceman, Ötzi, who died 5300 years ago in central Europe, was infected by an hpAsia2 strain with little or no African ancestry [4], suggesting that the admixture probably took place within the last few thousand years.

In Latin America, gastric cancer is a leading cause of cancer death, and some countries in the region have among the highest mortality rates worldwide [5]. However, the mortality rates vary in different geographic regions, both between neighboring countries and within nations [5,6]. Several studies have been performed comparing *H. pylori* ancestry in high- and low risk areas and have linked phylogeographic origin of the bacteria, as well as discordant origin of bacteria and host, to increased risk of gastric cancer development [2,7]. However, these studies have been performed using MLST analysis that, being based only on seven housekeeping genes, is limited in its resolution compared to whole-genome comparisons.

To investigate if American *H. pylori* strains have differentiated from those found in the Old World by mixture, genetic drift or natural selection, we combined hundreds of publicly available genomes with over hundred newly sequenced genomes of *H. pylori* sampled in Latin America (Mexico, Nicaragua, and Colombia), Europe, and Central Asia. We show that the American bacterial populations have undergone substantial evolution within 500 years and our results also suggest that *H. pylori* transmission biology has been as important as human migration in determining extant patterns of diversity.

## Results

We used the Chromopainter/fineSTRUCTURE pipeline [8,9] to assess the population structure within our global collection of isolates (n = 401, described in Supplementary table 1). Chromopainter infers the ancestry of the core genome of each isolate as a mosaic of DNA chunks from a “donor panel” of other genomes. We used one donor panel consisting only of Old world (European, African and Asian) isolates in order to investigate the origins of the New World strains; “*Old World painting*”, and one global donor panel of all isolates; “*Global painting*”, to investigate the relationships of the New World strains to each other. We excluded hspAmerind strains from the donor set because many of the strains have undergone post-Columbian admixture with other populations.

fineSTRUCTURE assigns individuals to populations with distinct ancestry profiles. We applied fineSTRUCTURE to the global painting to identify subpopulations in the dataset (Figure 1). In order to make display and reporting of the results tractable, we merged the most similar populations until 12 distinct populations remained, 5 of which are restricted to the New World. The “palette” of each strain, representing the proportion of chunks that come from each population in the donor panel, was determined for both the Old World (Figure 2A) and the Global (Figure 2B) painting. One of the twelve populations, hspMiscAmerica, contains isolates that are not particularly closely related to each other and should not be thought of as a coherent population (Figure 1).

**Figure 1.**
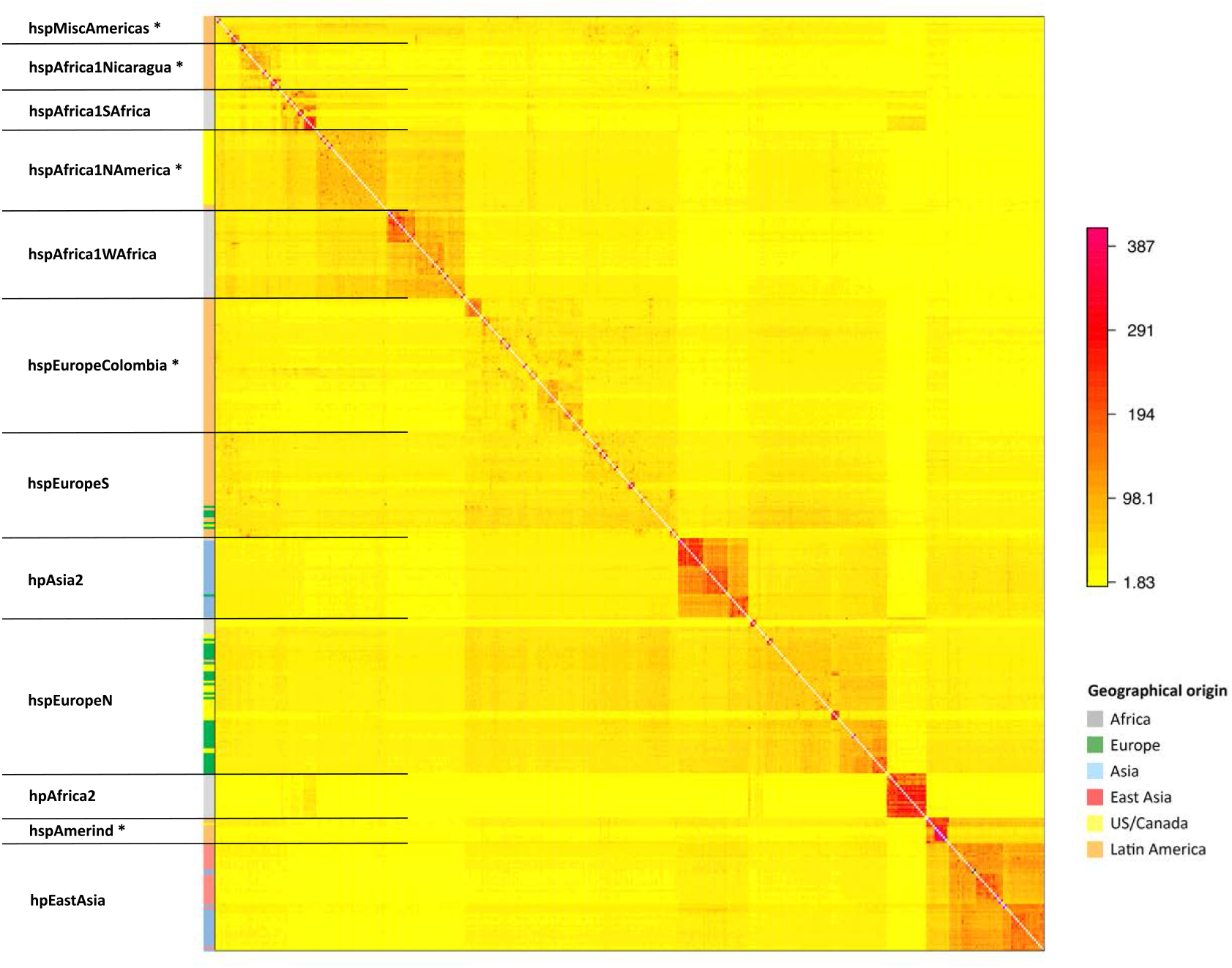
Population structure of global H. pylori strains. *The colour of each cell of the matrix indicates the expected number of DNA chunks imported from a donor genome (column) to a recipient genome (row). The boundaries between named populations are marked with lines, with New World populations marked with an asterisk. The colour bar on the left indicates the geographical locations where the strains were sampled.*

**Figure 2.**
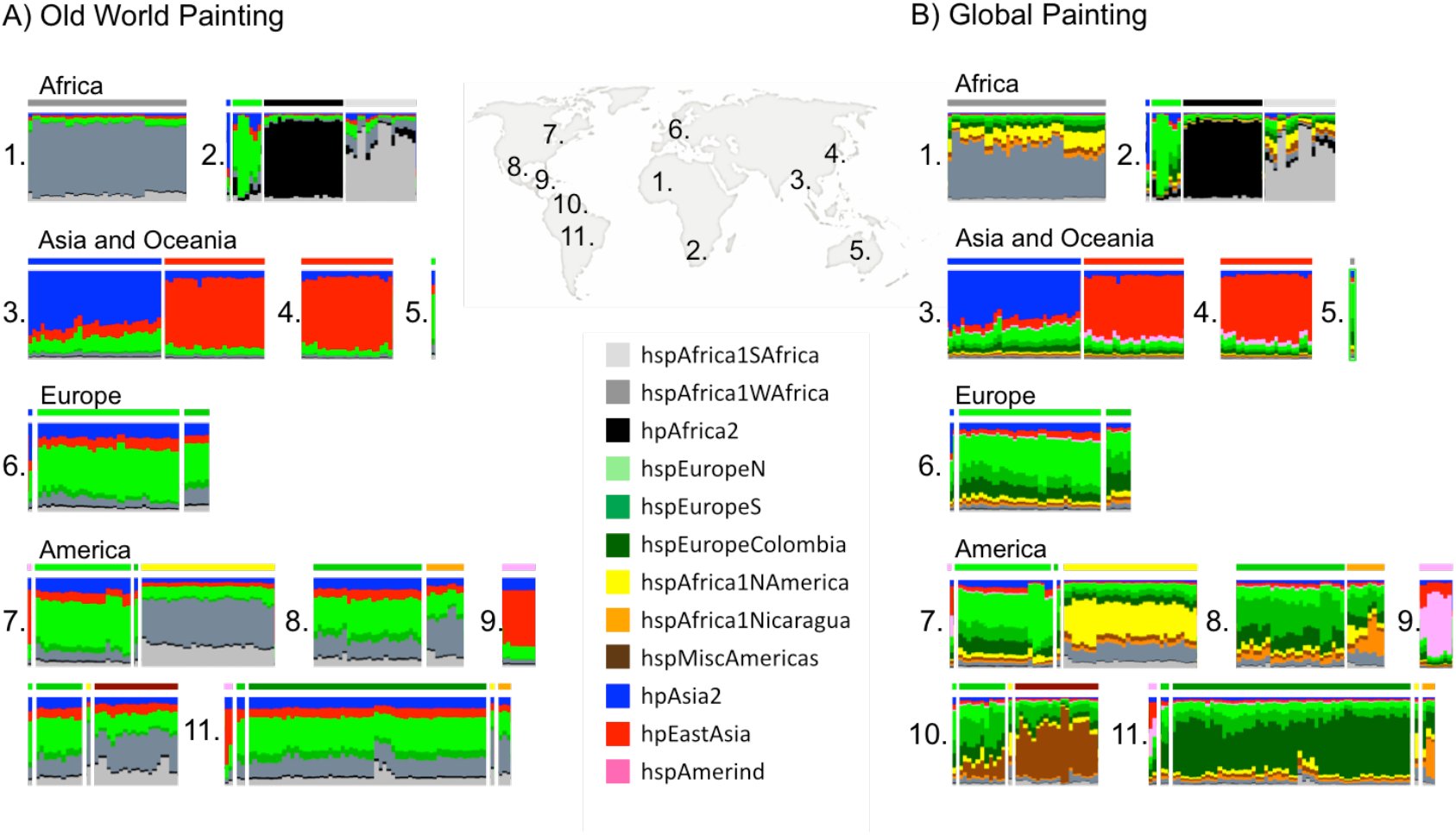
Ancestry of H. pylori inferred by chromosome painting. *Each vertical bar represents one isolate, which are ordered by geographical origin (1-11). 1: West Africa, 2: South Africa, 3: Central Asia, 4: East Asia, 5: Australia, 6: Europe, 7: US and Canada, 8: Mexico, 9: Central America, 10: Colombia, 11: Peruvian Amazon. The colour composition of each bar indicates each of the subpopulations’ contribution to the core genome of that isolate. A) Old world painting where only isolates from Old world areas (1-6 on map) have been used as donors in the chromosome painting. B) Global painting in which all populations have been used as donors.*

### Increased number of isolates reveals substructures in the Old World populations

Each of the 7 populations found in the Old World has been reported previously with the exception that, with the addition of the large number of isolates in this study, hpEurope isolates separated into two distinct groups, which we provisionally label hspEuropeN and hspEuropeS (Figure 1). Our geographical sampling within Europe is limited but this split is likely to reflect the previously observed North to South gene frequency cline [10,11], with the hspEuropeS isolates having a larger fraction of their palette from African populations and hspEuropeN having a higher proportion from HpAsia2. The other five populations, hpAfrica2, hspAfrica1SAfrica, hspAfrica1WAfrica, hpEAsia and hpAsia2 are highly distinct from each other, each receiving more than half of their palette from their own population in the Old World painting.

### Distinct subpopulations of mixed hpEurope and hpAfrica1 ancestry in American *H. pylori*

Among the isolates from the Americas, five additional subpopulations could be distinguished; Four have palettes consistent with being European/African hybrids, according to the Old World painting (Figure 2A). The population with the highest African ancestry is *hspAfrica1NAmerica*, isolated from 30 individuals in the US, one in Canada, one in Nicaragua, and one in Colombia, followed by *hspAfrica1Nicaragua*, which only contains isolates from Nicaragua; *hspMiscAmerica*, which consists of a number of strains of Mexican and Colombian origin; and *hspEuropeColombia*, which contains most of the Colombian isolates in our data set, and has a palette similar to hpEuropeS (Figure 1). The fifth population, hspAmerind, has a palette similar to hpEastAsia but with more hpEurope ancestry.

In our sample, several isolates from the Americas cluster within the two hpEurope subpopulations (Figure 1). The hpEurope strains from North America largely cluster with hspEuropeN while those from Central and Southern America cluster with hspEuropeS. There was also substantial variation in the proportion of the genomic palette stemming from hspAfrica1WAfrica and hspAfrica1SAfrica, both between and within New World populations. hspAfrica1WAfrica is the major African source in isolates from hspMiscAmerica, hspEuropeColombia as well as hspEuropeS while hspAfrica1SAfrica is a more important source for hspAfrica1NAmerica and hspAfrica1Nicaragua populations. A handful of isolates from both hspEuropeColombia and hspAfrica1Nicaragua populations have elevated hspAfrica1SAfrica proportions, consistent with recent genetic mixture (Figure 2A).

### The distinct New World subpopulations show evidence of both drift and mixture

To study the evolution of the New World populations after arrival in the Americas, we performed “global painting” where all the populations, both from the Old World and from the Americas, were included as donors (Figure 2B). In this painting, the strains from the New World populations receive most of their ancestry from within their own population, meaning that they are highly differentiated from the other New world populations as well as from all of the Old World isolates in the sample. The formation of differentiated populations in the Americas involves demographic bottlenecks but the New World populations have nucleotide diversity as high as or slightly higher than the Old World populations from which they evolved (Figure 3), presumably because the diversity lost in bottlenecks has been replaced by admixture.

**Figure 3.**
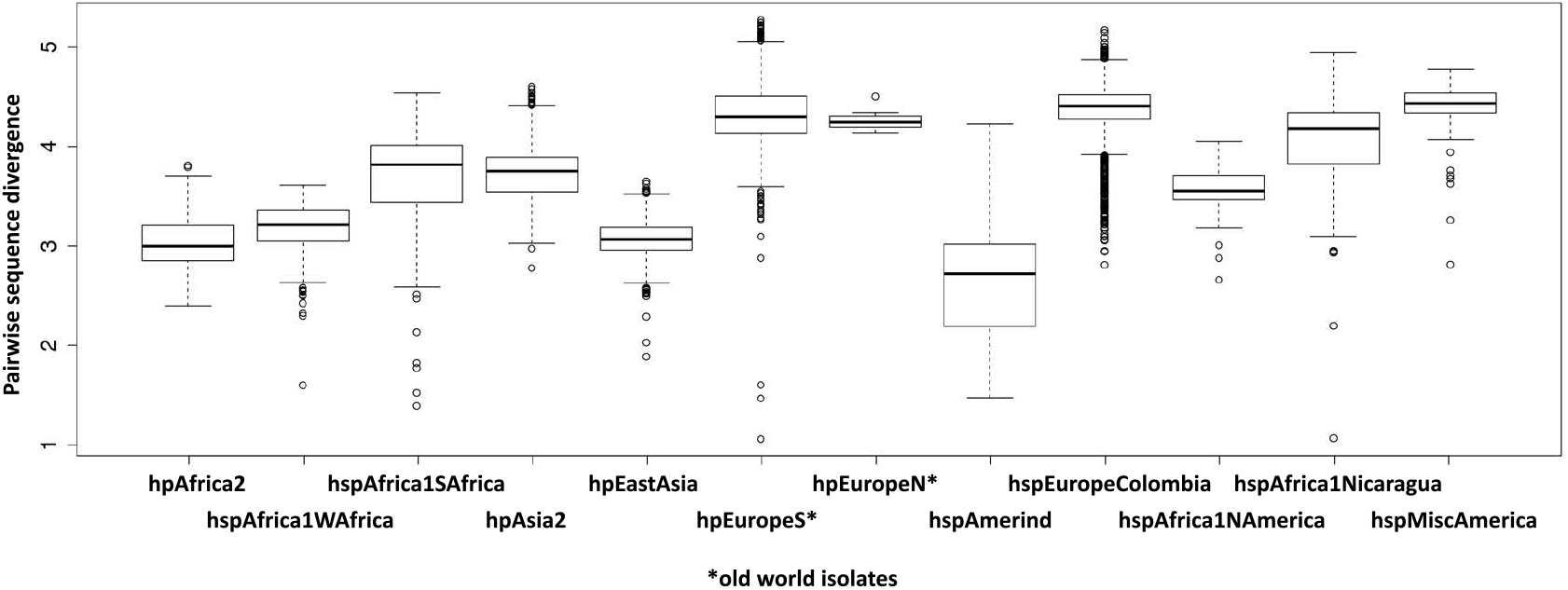
Pairwise sequence divergence within populations. *For the two hspEurope populations only the Old World isolates are included*.

Identifying the components of the ancestry of the New World populations that have undergone higher levels of drift provides insight into the process of differentiation. Drift is likely to be caused by the expansion of particular clones or lineages, for example, due to transmission bottlenecks. Specifically, we tabulated the proportion of “self-copying” for each Old World source, i.e. DNA that is assigned to that population in the Old world painting but is instead inferred to be derived from other members within the same population in the global painting, (Table 1). Bottlenecks allowing small numbers of clones to propagate will lead to high self-copying while diversity acquired by admixture is more likely to be copied from other populations, unless the admixture sources have themselves been subject to a strong bottleneck. For the hspAmerind population, the most drifted component is the component assigned to hpEastAsia, consistent with bottlenecks during the original colonization of the Americas by Native Americans, while in hspEuropeColombia, the European components are most drifted and for the other three populations, it is the African component. The difference is most pronounced for these last three populations, suggesting that African lineages may have undergone rapid demographic increases during their spread in the Americas and thus that they may have a transmission advantage.

**Table 1.** Proportion of ancestry assigned to each Old World population (columns) in the Old World painting that becomes self-copying within the global painting

|  | hspAfrica1SAfrica | hspAfrica1WAfrica | hspEuropeS | hspEuropeN | hpAsia2 | hpEastAsia |
| --- | --- | --- | --- | --- | --- | --- |
| hspEuropeColombia | 0.43 | 0.44 | 0.48 | 0.48 | 0.46 | 0.39 |
| hspAfrica1NAmerica | 0.42 | 0.41 | 0.34 | 0.27 | 0.27 | 0.29 |
| hspAfrica1Nicaragua | 0.61 | 0.66 | 0.43 | 0.45 | 0.47 | 0.50 |
| hspMiscAmerica | 0.11 | 0.21 | 0.06 | 0.06 | 0.06 | 0.06 |
| hspAmerind | 0.46 | 0.45 | 0.44 | 0.47 | 0.50 | 0.53 |

Aside from the isolation of hspAmerind strains from three countries and a single hspAfrica1NAmerica isolate from Colombian and Nicaraguan, there was no indication of sharing of ancestry between North, Central and South American gene pools. There is also no evidence from the palettes of hspAmerind having contributed DNA to any other New World strains. Amongst the Mexican isolates, a few hspMiscAmerica isolates have a substantial hspAfrica1NAmerica component but there is no sign of elevated ancestry from the Colombian or Nicaraguan populations.

The palettes provide evidence of genetic mixture between populations within countries. The hspEuropeS isolates from Nicaragua have more hspAfrica1Nicaragua in their palette than those from other locations, while Colombian isolates that are not assigned to the hspEuropeColombia have a higher ratio of hspEuropeColombia/hspEuropeS than found elsewhere, which is consistent with recent genetic exchange. Conversely, there is no evidence for elevated hspAfrica1NAmerica ancestry in hpEuropeN isolates from North America. The hspAfrica1NAmerica population has more hpEurope ancestry than hpAfrica1 isolates from Africa but there is little variation between strains, contrary to what would be expected if there was substantial ongoing gene flow.

### Several genes have ancestral origin distinct from the overall core ancestry

The spread of *H. pylori* populations in the Americas provides an opportunity to investigate adaptive introgression as the bacteria encountered new populations of humans, as well as novel diets and environmental conditions. This is of specific interest since *H. pylori* has an outstanding capacity for recombination between co-colonising strains [1,12]. We performed a scan of the core genome for genomic regions with enrichment of specific ancestry components. To this end, we painted the strains from each New World population, using Old World strains as donors and recorded whether the donor was European, African or Asian in origin. We found several genes where alleles showed significantly higher or lower ancestry from another Old world donor population than would be expected based on the overall ancestry of that isolate (Table 2). Among these were four genes that had unexpected ancestry in more than one of the New World populations. These were the genes encoding for TlpA (HP0009), HofC (HP0486), AlpB (HP0913), and FrpB4 (HP1512), which notably all are outer membrane proteins (Figure S1) and are enriched for Asian ancestry in at least one population.

**Table 2.**
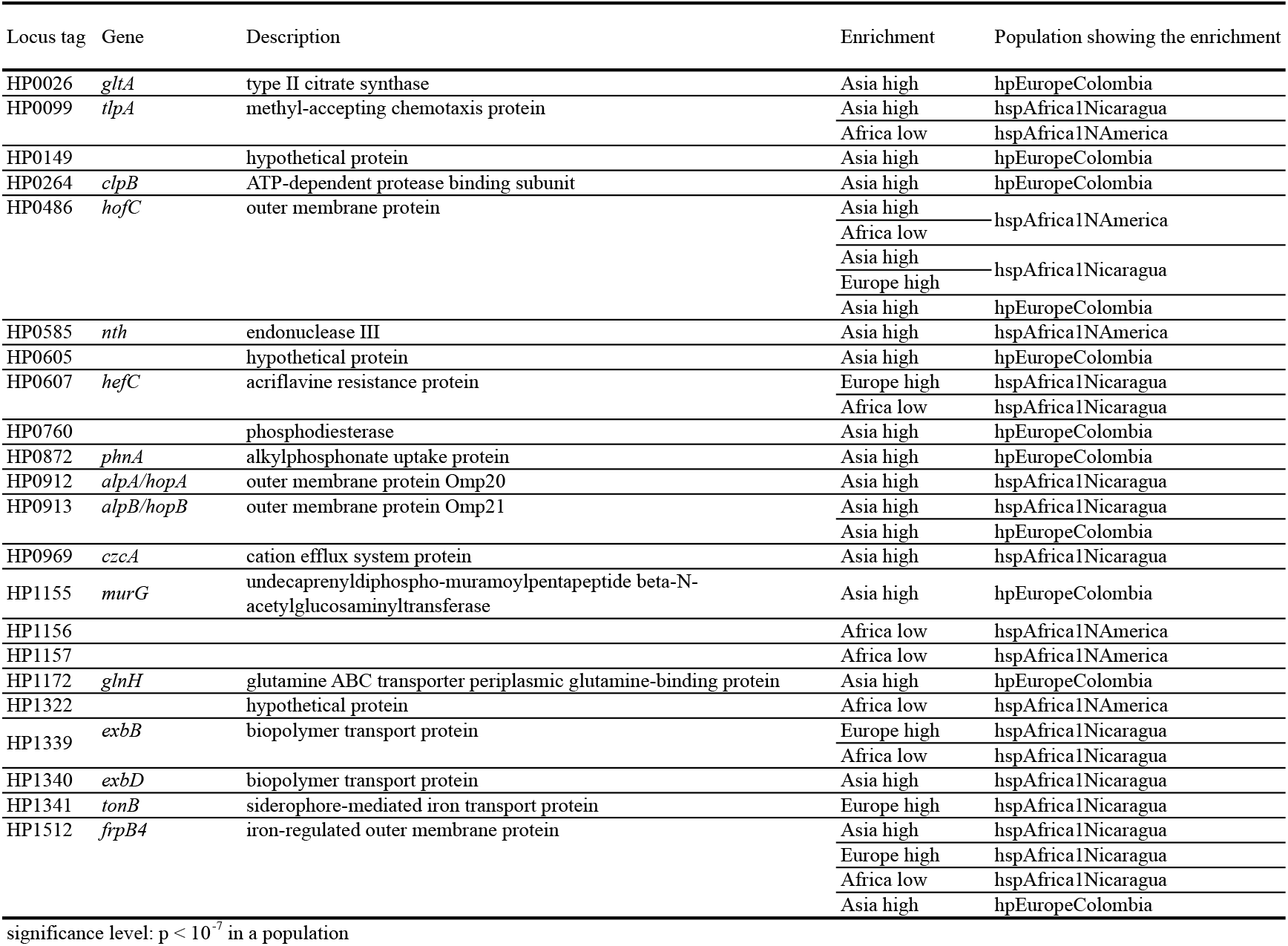
Genes carrying position(s) with enrichment of a specific ancestry components

To investigate if the enriched positions contributed to the overall phylogeny we constructed phylogenetic trees of the four genes. Especially interesting were the patterns found in *alpB* and *hofC* trees (Figure 4). For each gene at least one major separate clade of Latin American isolates could be observed, regardless of *H. pylori* population. Trees for *tlpA* and *frpB4* can be found in Figure S2.

**Figure 4.**
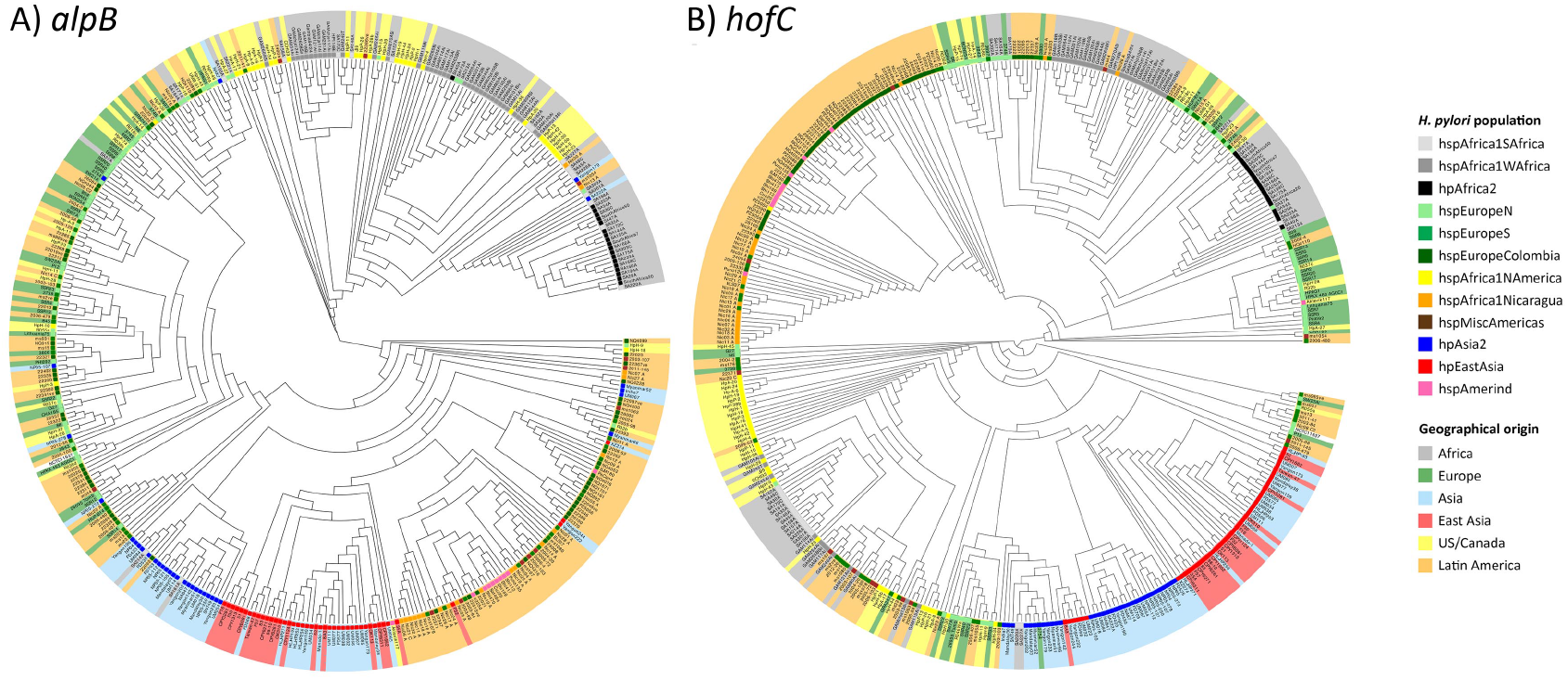
*Maximum likelihood phylogenetic trees of* alpB *and* hofC *genes*. *Leaves are shaded according to geographical origin and the* H. pylori *population assignment according to the FineSTRUCTURE analysis is marked at the base of each leaf*.

For *alpB* there are three major clusters; a predominantly Asian cluster including a majority of the Latin American strains, a predominantly European cluster, also with a number of Latin American strains, and an African cluster where isolates from Africa group together with isolates the hspAfrica1NAmerica. Notably, in the Asian group the Latin American isolates from multiple *H. pylori* populations cluster together while in the European group they are interspersed with the other isolates (Figure 4A). For *hofC* there is one clearly distinct South American clade, including all the Amerindian strains except for Aklavik117. The Other three main clades are dominated by either: (*i*) hspAfrica1WAfrica, hpAfrica2 and hspAfrica1NAmerica isolates; (*ii*) hspAfrica1SAfrica, European and US/Canadian hpEurope isolates or; (*iii*) Asian isolates, respectively (Figure 4B). Notably, for *hofC* the Mexican isolates did not group within the main South American clade but within clade *i* and *ii*.

Investigating the *hofC* gene alignment in more detail revealed that the sequence variation strongly contributing to the tree clade structure was nucleotides 826-840 of the gene, corresponding to 5 amino acids (Figure S3). Within this stretch, amino acids 277(V) and 279(T) were completely conserved in the alignment. The South American clade of the gene has a Glutamic acid instead of a Glycine at position 278, which is conserved among all the other strains. At position 280, all other isolates have Leucine, while isolates in the South American clade have Asparagine or Aspartic Acid (Table 3). This motif has spread to a large proportion of isolates in all of the populations found in South America, suggesting they confer an adaptive advantage.

**Table 3.** Amino acids 276-280 of HofC

|  | 276 | 277 | 278 | 279 | 280 |
| --- | --- | --- | --- | --- | --- |
| South American clade | E/Q | V | E | T | D/N |
| Other isolates | Q/M/E | V | G | T | L |

### Accessory genome analysis shows similar but not identical ancestral patterns to the core genome

Our collection of multiple genomes from each population allowed us to examine patterns of gene presence and absence. A neighbour-joining tree based on gene sharing distance between isolates largely recovered the populations and sub-populations identified based on core genome sequence, but with distinct clusters for cagPAI positive and cagPAI negative isolates (Figure 5A).

**Figure 5.**
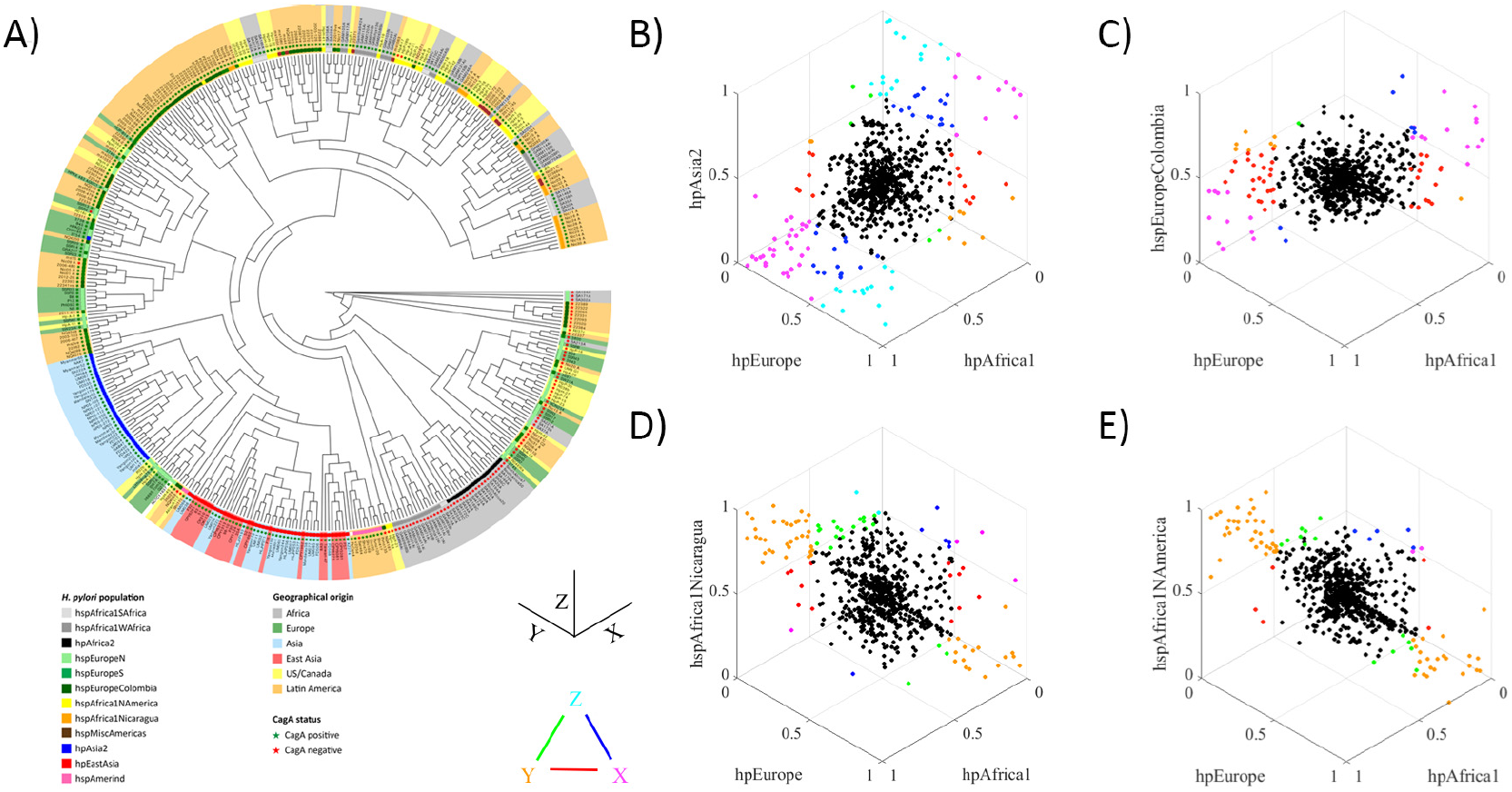
Accessory genome analyses. A) Neighbour-joining tree based on gene sharing distance. The colours are the same as those in Figure 4 and are indicating geographical origin (leaf shading) and population assignment to according to the FineSTRUCTURE analysis (colour at the base of each leaf). Green stars mark strains with the Cag pathogenicity island (CagPAI) and red stars are strains without a CagPAI.
B-E) Frequency of genes in the accessory genome compared between groups. The graphs are orientated such that genes with identical frequencies in all three populations appear at the centre of the plot. Genes with large frequency differences between populations are labelled in colours, according to the triangular legend. Colours shown on the vertices indicate genes that differ substantially between one population and the other two (according to the criteria that X is considered substantially bigger than Y if X – Y >= 0.5, X >= 0.5 and Y < 0.1 or X > 0.9 and Y <= 0.5), while colours on the edges indicate genes where the two populations on the vertices differ substantially in frequency, with the third population having an intermediate frequency. B) Comparison between Old world populations hpEurope, hpAsia2 and hpAfrica1, C) Comparison of hspEuropeColombia to hpEurope and hpAfrica, D) Comparison of hspAfrica1Nicaragua to hpEurope and hpAfrica, E) Comparison of hspAfrica1NAmerica to hpEurope and hpAfrica.

It has been previously shown that for the core genome, hpEurope bacteria are hybrids between hpAsia2 and hpNEAfrica (which is related to hpAfrica1), with higher hpAsia2 ancestry proportions in Northern Europe [10,13]. This could also be observed in our analysis, where the hpEurope population has a profile that is intermediate between that of hpAsia2 and hpAfrica1 (Figure 5B).

For the New World populations, the patterns for individual genes showed that hspAfrica1Nicaragua and hspAfrica1NAmerica have pan genomes that are more similar to those of hpAfrica1 than hpEurope (Figure 5D, E), while hspEuropeColombia has a profile that is intermediate between Africa1 and European isolates (Figure 5C). In both hspAfrica1Nicaragua and hspEuropeColombia however, the African contribution to the accessory genome is slightly higher than in the core genome painting. This pattern might be explained by recent admixture in America, but it might also reflect differences between the population of bacteria that arrived in America and the profile of the hpEurope isolates in our sample, which are mostly from Northern Europe.

## Discussion

Millions of people from diverse geographical and ethnic backgrounds have migrated from the Old World to the Americas in the last 500 years and it is likely that a majority carried *H. pylori*. Transmission between ethnicities and DNA exchange between strains might be expected to scramble the relationship between bacterial and human ancestry at the individual level, but in the absence of selection or bottlenecks, overall *H. pylori* ancestry should largely recapitulate the ancestry found in humans [10,13,14]. Consistent with this expectation, we find diverse populations of hpEurope bacteria in Northern and Latin America, with chromosome painting profiles comparable to those found in European isolates. We find a broad North-South divide amongst hpEurope isolates, both in the New and Old World, with higher relatedness to hpAfrica1 DNA in the southern populations. This is consistent with the gene frequency cline already observed in Europe and known differences in the colonization history of North and South America [15].

However, *H. pylori* genomic variation does not necessarily recapitulate patterns found in humans. The Americas constituted both a new physical and dietary environment and a new ethnic mix of hosts. Particular bacterial lineages may have had, or acquired, traits that adapted them to these new conditions. In extreme cases, human migrations that have little or no effect on human ancestry might precipitate substantial changes in *H. pylori* populations. For example, hspAmerind strains are rare even in populations with substantial Native American ancestry [2]. This suggests that after more than 10,000 years of independent evolution, resident *H. pylori* lineages were poorly equipped to compete with incoming lineages or with changes in the environment caused by the new settlers. We also found evidence of substantial differentiation of New World *H. pylori* populations from their ancestors, which suggests that there have been bottlenecks with particular lineages contributing disproportionately to extant populations. These bottlenecks have most strongly affected African components of ancestry (Table 1), suggesting that bacteria of African origin may have been particularly effective in colonizing the new continent.

We identified three differentiated populations in the Americas, in addition to hspAmerind. The hspAfrica1NAmerica population includes the non-European isolates from the US, also found in single Canadian, Colombian and Nicaraguan isolates. This population has an ancestry profile consistent with it being a mix of West African, South African and European sources. However, our global chromosome painting results (Figure 2B) show that within genomic regions of African origin, the DNA of hspAfrica1NAmerica is distinct from that found in modern Gambian and South African populations. Differentiation at the DNA sequence level is also found in the hspEuropeColombia and hspAfrica1Nicaragua populations, whose gene pools are distinct from each other and from those in Mexico and Europe.

The three larger groups of samples, from Mexico (Mexico City), Nicaragua (Managua) and Colombia (Bogotá) respectively, were all collected at hospitals that are tertiary referral centres for endoscopy with large catchment areas, while all but one of the US isolates came from a hospital in Cleveland, a cosmopolitan city. Therefore, our findings likely reflect broad patterns of diversity within large geographic regions. Within our sample, there are regional differences in the proportions of European, African and Amerind ancestry and wider sampling might have differentiated the picture further. Nevertheless, the distinct patterns of *H. pylori* ancestry in the four countries indicate that recent population movements have been strongly influenced by national boundaries.

*H. pylori* can undergo high levels of recombination during mixed infection. Over time, this might lead to bacteria acquiring an ancestry profile that reflects their local gene pool rather than their continent of origin. Recombination has not proceeded this far anywhere in the Americas and multiple populations with distinct ancestry profiles are found in most locations. hspAmerind strains have not contributed substantially to the ancestry of bacteria from any other population, but do appear to have acquired hpEurope DNA themselves. In Nicaragua and Columbia, recombination has transmitted distinctive DNA between populations, e.g. the brown shaded component in the hpEuropeS isolates from Nicaragua (Figure 2B), leading to what can informally be thought of as a national signature in the *H. pylori* DNA. There is no equivalent signal of hspAfrica1NAmerica DNA amongst the hpEurope bacteria from the US, indicating that recombination between these populations has been less extensive, and there is also no evidence within our sample of a distinctive population of hpEurope bacteria evolving within the US. Similar patterns of higher admixture in African American and Hispanic American individuals than in American individuals of European descent have been observed also on human genomic level [15].

The differences in the extent of admixture in the New World populations can have several explanations including differences in dates of colonization and extent of European and African influx/admixture in Latin America compared to the US. Another important factor can be the prevalence of infection in different areas. The prevalence of *H. pylori* infection remains high in Latin American countries, ranging from 70,1 % to 84,7 % of adults in a recent multi-country study [16]. In the US, the prevalence has been declining from high levels and according to data from the end of the 1990’s, is around 32,5 % [17]. The prevalence was different between the ethnic groups: 52.,7 % in non-Hispanic blacks; 61,6 % in Mexican Americans and; 26,2 % in non-Hispanic whites [17]. High prevalence likely entails higher occurrence of horizontal transmission and mixed infections and thus the possibility of recombination between distantly related strains [18] [19].

Our sample of Old World sources is incomplete, both in Africa and Europe, and therefore it is likely that Old World sub-populations exist that are more closely related to the New World populations than those in our sample, for example, in Spain. Sampling limitations also make it unclear how much of the extensive mixture between African and European DNA observed in many Central and Southern American isolates actually took place in the Americas. Nevertheless, it is difficult to explain the local affinities within the diverse gene pools in both Nicaragua and Colombia, except by local genetic exchange. The hpAfrica1NAmerica isolates are homogeneous in their ancestry profile, suggesting that they also form a distinct gene pool that has acquired its characteristics through substantial evolution within the USA.

The approximate panmixia within hspAfrica1NAmerica isolates (e.g. the relatively even shading of the within-population coancestry in Figure 1) is difficult to reconcile with the low levels of genetic exchange observed with hpEurope isolates from the US. Since it has been shown that *H. pylori* from the same population (hpEastAsia) can exchange 10% of their genome during a single 4-year mixed infection in human [21], the ancestral pattern in US *H. pylori* implies barriers to recombination between the two populations. Such barriers may be the result of ethnic segregation and thus less diverse co-infections, of differential uptake or incorporation of DNA from different populations, or of efficient competitive exclusion of bacteria from one population by bacteria from the other within individual stomachs.

In the New World populations, four genes encoding for outer membrane proteins have sequence with ancestry that differed from that inferred for the overall core genome. Interestingly, several of these variants were common for Latin American isolates regardless of which ancestral population they belonged to. AlpB is an adhesin binding to laminins in the extracellular matrix [22] that is present in all *H. pylori* strains [23]. Together with AlpA, it is required for colonization in experimental models and for efficient adhesion to gastric epithelial cells [24]. The HofC protein is also required for *H. pylori* colonization in mice and gerbils [25,26] is not well characterized and little is known about its function. TlpA and FrpB4 are important in the bacterial adaptation to variation in the microenvironment. TlpA is required for arginine and bicarbonate chemotaxis [27], and FrpB4 is regulated by the levels of nickel, a micronutrient essential for *H. pylori* survival, growth and expression of virulence factors in the human stomach [28-30].

The enrichment pattern in *hofC* in a high number of the South American isolates was largely explained by two amino acids in position 278 and 280 of the 528 amino acid protein. These two variants were found in all the South American Amerindian strains as well as almost all of the hspAfrica1Nicaragua and a majority of hspEuropeColombia strains together with strains from Peru and El Salvador. No Mexican strains were found in this clade. Since the HofC protein structure and function are not characterised in detail, it is unknown how these alleles contribute to the function or specificity of the protein. Nevertheless, the very pronounced enrichment pattern, as well as that in the other genes, is consistent with the New World *H. pylori* having adapted to their respective human populations, allowing certain traits to propagate relative to the overall genetic background. This could be important in understanding the differences in pathogenicity in different areas and different host/bacterial interactions, suggesting a need for further investigation of the function of these proteins.

Our analysis of the accessory genome shows that *H. pylori* gene content, as well as nucleotide composition, is mixed during admixture between host populations. For example, the gene content of hpEurope is intermediate between that of hpAfrica1 and hpAsia2, but with substantial variation that may reflect the large time that has elapsed since admixture. hpEuropeColombia is more African in genome content than the average hpEurope bacteria from Europe, as would be expected because its higher African ancestry at the nucleotide level. However, the genome content of strains from the hpAfrica1Nicaragua population is more African than would be expected given its substantial co-ancestry with hpEurope within the core genome. This observation is concordant with recent observations showing that restriction modification inhibits non homologous but not homologous recombination [31], suggesting that core genome ancestry may mix more readily between populations than accessory elements if restriction modification is an important barrier to exchange.

Our results on the population structure in the Americas sheds new light on the relationship between human migration and *H. pylori* diversity. In particular, we show that at least during human population upheavals, evolution within geographic locations is far more dynamic than the broad correlation with human genetic variation would suggest and that novel subpopulations can arise by a combination of genetic drift and admixture within hundreds of years.

## Acknowledgements

We thank all the researchers worldwide that have whole-genome sequenced *Helicobacter pylori* isolates and made their data available to us, either by personal connections or by making the data publicly available. This work was supported by the Grants-in-Aid for Scientific Research from the MEXT (Ministry of Education Science, Sports and Culture) (15K21554 to K.Y.) and a Wellcome Trust Career Development fellowship and funding under the MRC-CLIMB initiative (to S. K. S. and D. F.). D.J.L. is funded by Wellcome Trust and Royal Society grant WT104125AIA. The computational calculations were done at HPC Wales and at the Human Genome Center at the Institute of Medical Science (the University of Tokyo).

## Materials and Methods

### Helicobacter pylori whole genome sequencing data

We used both publicly available and newly sequenced genomes of *H. pylori* isolates, 401 in total (Supplementary Table S1). Nicaraguan isolates were collected at Hospital Escuela Antonio Lenin Fonseca (HEALF) in Managua, within the international collaboration “Immunological Biomarkers in Gastric Cancer development” and previously described in [32]. Colombian isolates that are not previously described were collected at the Oncology hospital (INCAN) in Bogota, and the Mexican isolates were collected at the Oncology and General Hospital in Mexico City. All three hospitals are tertiary referral centres for endoscopy and patients may thus come from other locations within the countries. For the cases we had more detailed data on the origin of the individuals, this is noted in Supplementary Table S1.

The publicly available Colombian and North American genomes were those reported in a preceding studies, i.e [20,33,34].

### Data preparation

All of the genome sequences were imported into the Bacterial Isolate Genome sequence database (BIGSdb) [35]. After this, a gene-by-gene alignment was performed using CDS sequences of the *H. pylori* 26695 strains as reference, and the alignments were exported from the database. The exported file will be available at the public data repository Dryad (http://datadryad.org/) on publication. We conducted SNP calling for each alignment, and imputation for polymorphic sites with missing frequency < 1% using BEAGLE [36] as our preceding study [37]. We combined in total 401350 SNPs in 1232 genes while preserving information of SNP positions in the reference genome, to prepare genome-wide haplotype data.

### Population structure analysis

We inferred population structure among the strains from the genome-wide haplotype data by using the chromosome painting and fineSTRUCTURE [8], according to a procedure of our preceding study that applied them to *H. pylori* genomes [9]. We used ChromoPainter (version 0.04) and fineSTRUCTURE (version 0.02) software.

### Stratified chromosome painting

We conducted two types of chromosome painting; “Old World chromosome painting” using only Old world isolates as donors, and “Global chromosome painting” in which each isolate is painted using all of the others. For this purpose, we used ChromoPainterV2 software [8].

To identify genomic regions with enrichment of unexpected ancestry components in the New World populations hspAfrica1NAmerica, hspAfrica1Nicaragua, and hspEuropeColombia, we conducted a novel statistical test for each of the 401350 SNPs. This was done using the Old world strains as donors, grouped into African, Asian and European geographic origin respectively.

We aim to count the number of recipient haplotypes from a certain donor population at each SNP. However, we do not observe whether a recipient *i* uses a particular donor population a, but instead the probability that it does at each locus *l*.

The distribution of the total number of isolates at locus *l* from donor population *a* is ∼ Poisson-Binomial(*p_lia_*). If we let the genome-wide painting probability be 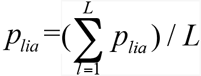, then the distribution expected under the null that there is no local structure to the painting donors is ∼ Poisson-Binomial (*p_ia_*). We therefore report the p-values to test whether locus *i* has significantly enriched for donor *a* (and likewise to test for de-enrichment). We used P<10^−7^ as a significance level, which corresponds to P<0.05 after Bonferroni correction.

Because a) the variance of a Poisson-Binomial is highest when is close to 0.5, and b) the distribution is discrete, this statistic has less power to detect high ancestry contributions from components that have high genome-wide ancestry, especially when sample size is small. In practice this has limited our power to detect regions that have an excess of African ancestry.

### Phylogenetic analysis of genes with enriched ancestry

Multiple alignments of the genes were performed using MUSCLE [38] and the alignment manually inspected to remove sequences with incomplete coverage before a PhyML maximum likelihood tree was created using the SeaView software [39]. All trees were visualized using Evolview [40].

### Analysis of gene presence/absence and accessory genome

A pan-genome was constructed with all loci present in at least one of our 401 strains to examine presence/absence of all *H. pylori* genes. This pan-genome list of 2462 genes was used as queries of BLASTN against each genome analysed in this study through the BIGSdb Genome Comparator pipeline [35]. Gene presence was judged by a BLASTN match of ≥ 70% identity over ≥ 50% of the locus length [41].

### Accessory Presence/Absence Tree

The Genome Comparator Output matrix obtained with BIGSdb was used to build a distance matrix (MATLAB R2015a, The MathWorks, Inc., Natick, Massachusetts, United States). A tree was obtained using SplitsTree4 [42] and was visualised with Evolview [40].

## Supplementary Figures

**Figure S1.**
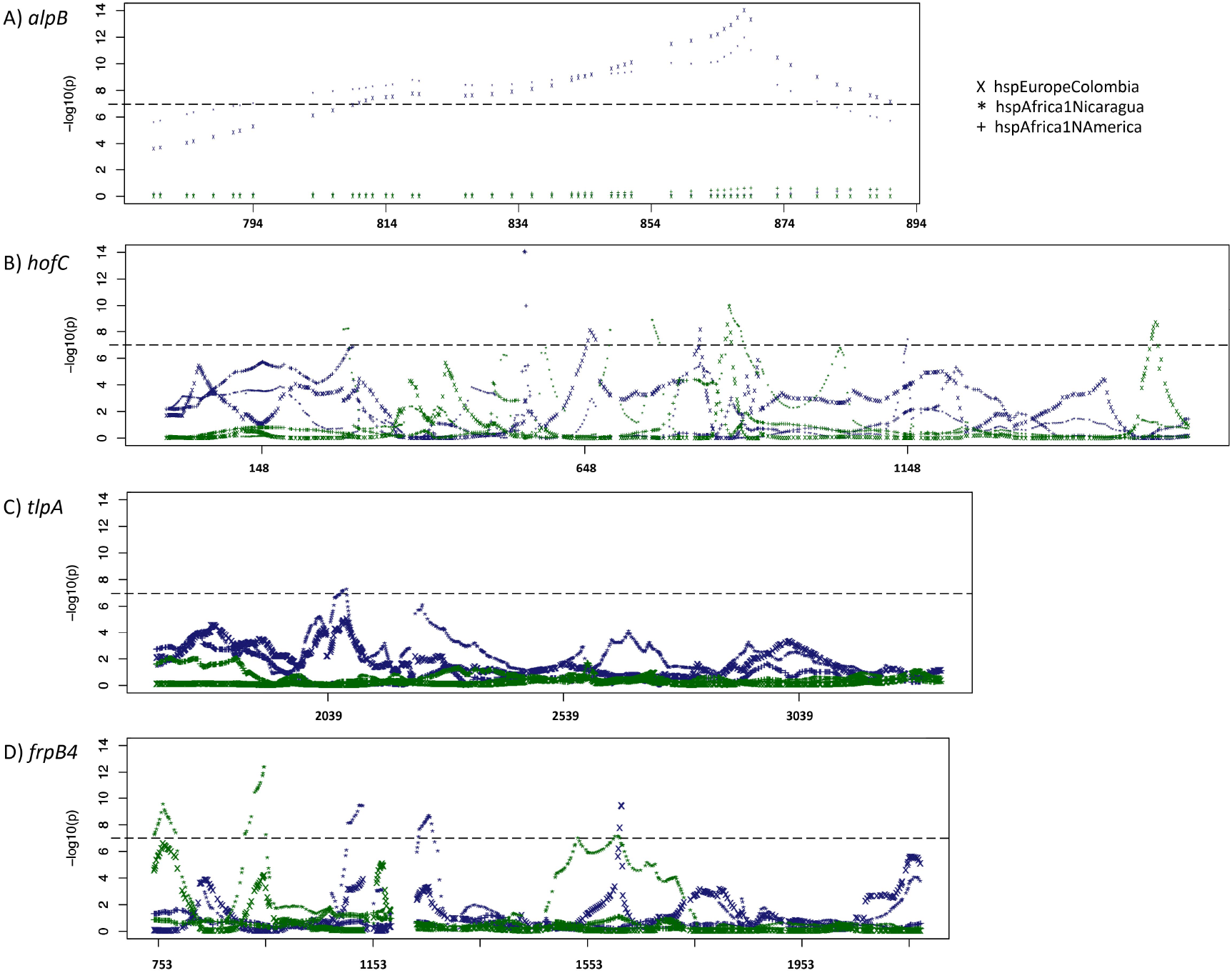
P-values for enrichment of European and Asian ancestry over genes. *Each dot corresponds to a polymorphic site that was tested statistically. The four genes in Table 1 satisfying significance level p < 10^−7^(p < 0.05 after Bonferroni correction) in more than one of the New World populations are shown. Blue symbols indicate the strength of statistical evidence for Asian enrichment and green European enrichment. A)* alpB, *B)* hofC, *C)* frpB4, *and D)* tlpA.

**Figure S2.**
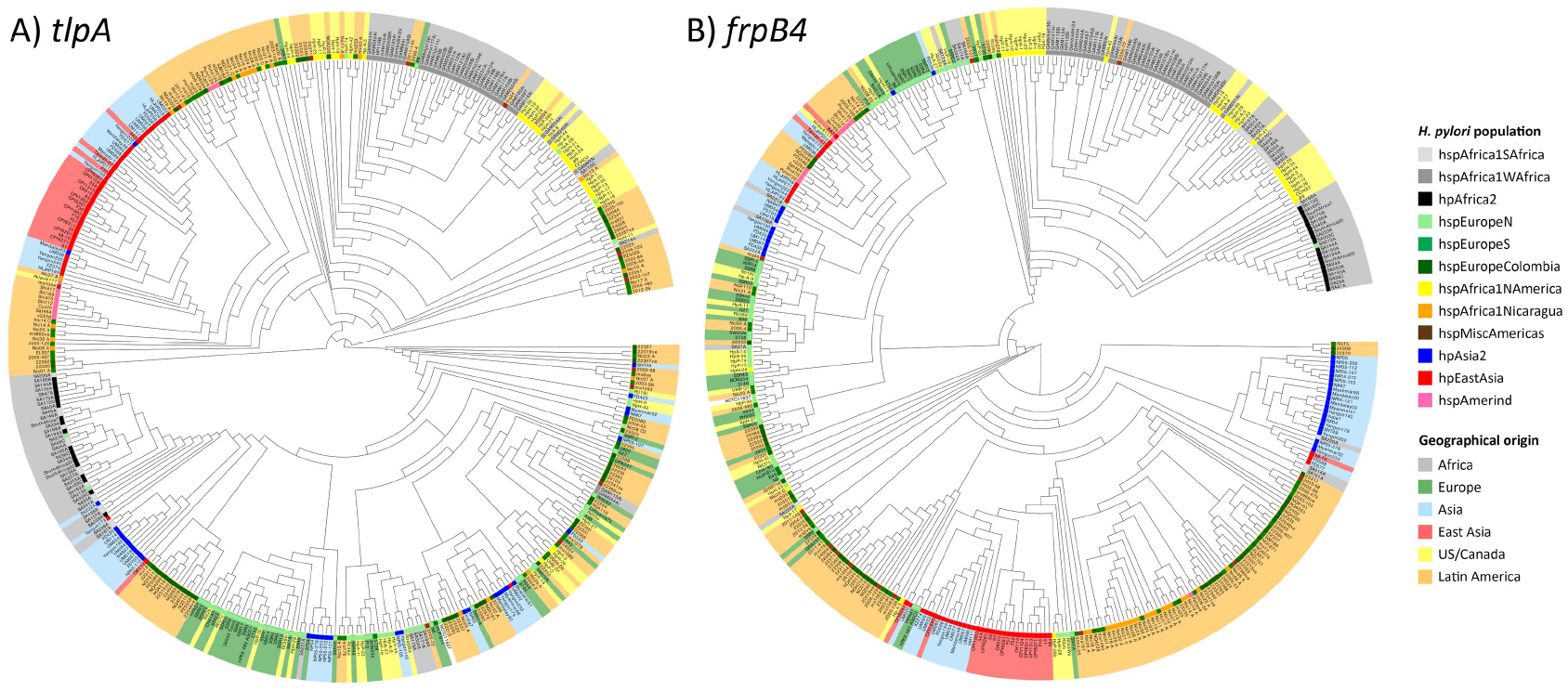
*Maximum likelihood phylogenetic trees of* tlpA *and* frpB4 *genes*. *Leaves are shaded according to geographical origin and the* H. pylori *population assignment to according to the FineSTRUCTURE analysis is marked at the base of each leaf.*

**Figure S3.**
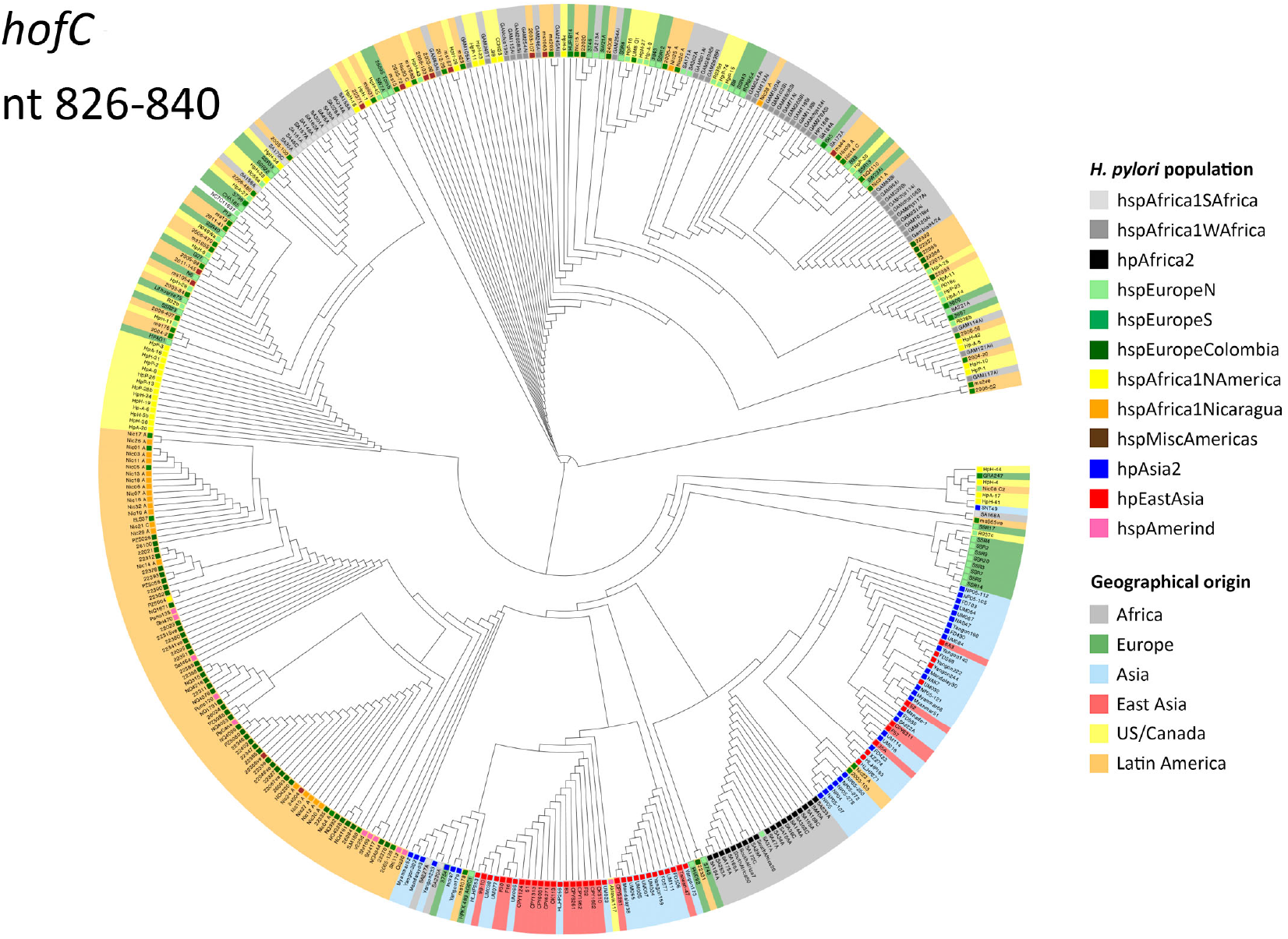
*Maximum likelihood trees for* hofC, *nt 826-840*. *Leaves are shaded according to geographical origin and the* H. pylori *population assignment to according to the FineSTRUCTURE analysis is marked at the base of each leaf.*

